# Cytokine-mediated activation of exhausted T cells

**DOI:** 10.64898/2026.09.28.755149

**Authors:** Eva Goljat, Rahel Daetwyler, Nima Yassini, Nikolas Rakebrandt, Zhaorong Li, Francisco Javier Quintana, Nicole Joller

## Abstract

1.

CD8 ⁺ T cell exhaustion during chronic infection and cancer is marked by impaired effector function and limited responsiveness to stimulation. While memory CD8⁺ T cells can be activated TCR-independently by IL - 12 and IL - 18, it remains unclear whether exhausted cells, despite diminished TCR responsiveness, retain this cytokine-driven effector capacity. Here, we show that IL - 12 and IL - 18 stimulation induce robust IFNγ production in exhausted CD8 ⁺ T cells from chronic LCMV Cl13 infection, exceeding responses to antigen-specific stimulation. This activation does not result in upregulation of Tox, suggesting engagement of transcriptional circuits distinct from those promoting exhaustion. Single-cell transcriptomics revealed an innate acting T cell (T_IA_)-specific gene signature enriched within exhausted populations in chronic infection and tumor-infiltrating lymphocytes, particularly at the interface of intermediate and terminal exhaustion. Using heterologous Legionella pneumophila infection, we confirmed in vivo bystander responsiveness of exhausted CD8 ⁺ T cells, reducing bacterial burden, highlighting their functional capacity. Our findings demonstrate that exhausted CD8 ⁺ T cells retain inducible effector capacity through IL - 12 and IL - 18 stimulation, providing new insights into possible routes to activate exhausted T cells.

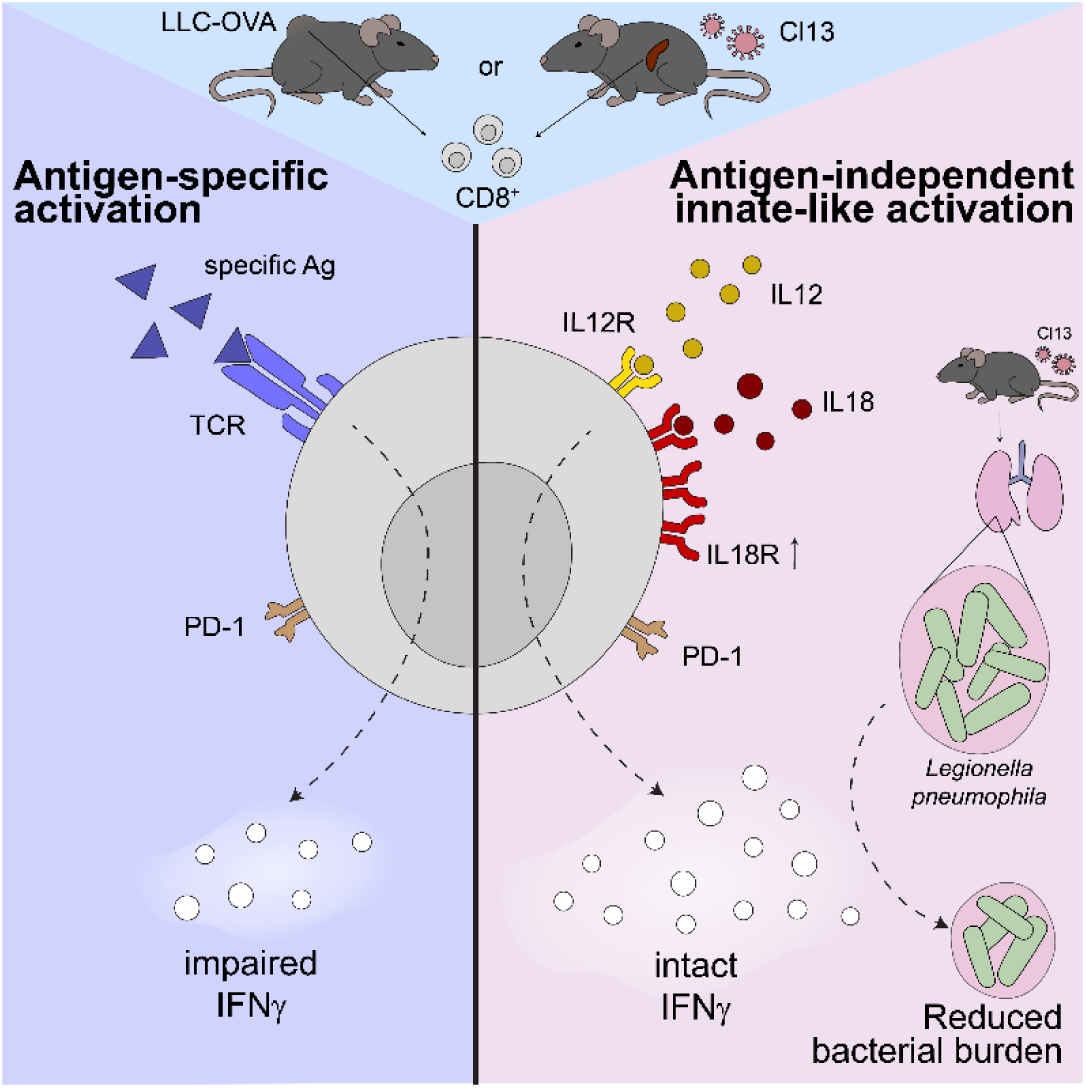

## 2. INTRODUCTION

CD8^+^ T cells are integral to the control of intracellular pathogens and tumors, due to their ability to recognize and eliminate infected or transformed cells, and to produce inflammatory cytokines such as IFNγ. Upon acute challenge, they differentiate into effector cells that clear the pathogen and give rise to long-lived memory cells (Kaech & Cui, 2012). However, during chronic infections such as lymphocytic choriomeningitis virus (LMCV) Clone 13 (Cl13) infection, persistent antigen exposure and ongoing inflammation drive CD8^+^ T cells towards a distinct state of T cell exhaustion (Wherry et al, 2007; Richter et al, 2012). Exhausted T cells are characterized by expression of inhibitory receptors, most notably PD-1 as well as a progressive decline in effector cytokine production, proliferative capacity, and cytotoxicity (Wherry et al, 2007; Blackburn et al, 2009). More recent studies have shown that exhaustion is a heterogeneous state, with some CD8 ⁺ T cells maintaining partial functionality and transcriptional features reminiscent of memory cells (Utzschneider et al., 2016; Zehn et al., 2022).

In parallel to findings on T cell exhaustion, a growing body of work in the past has highlighted the existence of bystander-activated or innate-like CD8 ⁺ T cells—cells that can mount effector responses in the absence of cognate antigen recognition, and remain an active area of investigation (Selin et al, 1994; Kim & Shin, 2019; Balint et al, 2024; Chu et al, 2013; Maurice et al, 2026). These cells, typically within memory compartments, can produce IFNγ and express cytotoxic molecules upon stimulation with cytokines alone. A seminal study by Freeman et al. demonstrated that IL - 12 and IL- 18 synergistically activate murine memory CD8 ⁺ T cells to rapidly secrete IFNγ, independent of TCR stimulation (Freeman et al, 2012). Innate-like bystander activation of memory CD8 ⁺ T cells via IL-12 and IL-18 has been described across several models and was summarized in recent reviews (Yosri et al, 2024). This innate-like responsiveness has been proposed to enable early protective responses and to bridge innate and adaptive immunity in contexts where TCR engagement is limited or delayed. This concept has since also been expanded to CD4 ⁺ memory T cells, where IL - 12 and IL - 18 alone are sufficient to trigger robust cytokine production in the absence of antigen (Guoa et al, 2009; Kawabe et al, 2017; Rakebrandt et al, 2024), supporting the broader relevance of this innate-like activation pathway across T cell subsets.

While bystander activation via IL - 12 and IL - 18 is well described in memory CD8 ⁺ T cells (Freeman et al, 2012; Balint et al, 2024), its role in chronically stimulated or exhausted contexts remains poorly understood. Tumor-infiltrating lymphocytes (TILs) are also frequently characterized by dysfunctional or exhausted states (McLane et al, 2019; Jiang et al, 2015; Schietinger et al, 2016). Immune checkpoint blockade has demonstrated that these cells can, under certain conditions, be functionally reinvigorated (Pauken et al, 2016); however, therapeutic responses are variable and often incomplete, indicating that alternative or complementary modes of activation may exist (Philip et al, 2017). We therefore asked whether cytokine-mediated, antigen-independent activation – reminiscent of bystander or innate-like CD8^+^ T cell responses – could represent an additional mechanism by which exhausted T cells might acquire effector functions. Moreover, direct comparisons of their functional output and transcriptional profiles are limited.

In this study, we investigated whether exhausted CD8 ⁺ T cells retain the capacity for an innate-like activation via IL - 12 and IL - 18. We show that cytokine stimulation elicits robust IFNγ production in exhausted CD8 ⁺ T cells, exceeding that of antigen-specific responses. Transcriptional profiling revealed similarities between exhausted and innate-like memory CD8 ⁺ T cells. Moreover, using heterologous bacterial infection, we demonstrate that this bystander responsiveness translates into functional protection in vivo. Together, these findings reveal that exhausted CD8 ⁺ T cells preserve inducible effector capacity through cytokine stimulation.

## 3. RESULTS

### 3.1. Exhausted T cells can be activated in a bystander manner

We first tested whether T cells from an exhausted setting, such as LCMV Cl13 infection, can be activated in a bystander manner in vitro to produce IFNγ. We compared the responsiveness of splenocytes from Cl13-infected mice at the chronic stage of infection (day 25–30), from LCMV WE memory mice (>day 40 post-acute infection) as a positive control, and from naïve mice as a negative control in various stimulation conditions (Fig. 1A). To our surprise, cytokine stimulation of CD8^+^ T cells from LCMV Cl13 infected mice induced IFNγ production that was comparable to anti-CD3 (αCD3) stimulation but lower than that observed upon stimulation with PMA/ionomycin (Fig. 1B). Cytokine stimulation even resulted in a greater effector response than antigen-specific stimulation with the immunodominant LCMV gp33 peptide (Fig. 1B and C), while the response to gp33 stimulation was markedly reduced compared with CD8⁺ T cells from acutely infected mice (Fig. EV1A), confirming their impaired antigen-specific responsiveness. This was also true when focusing on the exhausted CD8⁺ PD-1 ⁺ population (Fig. 1C and Fig. EV1B), indicating that, in exhausted T cells, IL-12 and IL-18 stimulation induces a stronger IFNγ response than antigen-specific stimulation. As expected, gp33-tetramer ⁺ LCMV-specific exhausted PD-1^hi^CD8⁺ T cells showed high expression of Tox, a key regulator and marker of T cell exhaustion (Yao et al, 2019; Khan et al, 2019)(Fig. EV1C, D), which. We did not observe any shifts between PD-1 intermediate (PD-1^int^) or high (PD-1^hi^) populations under the different stimulation conditions (Fig. EV1E) indicating that different types of stimulation do not lead to shifts in PD-1 expression. Importantly, cytokine stimulation of exhausted CD8^+^ T cells did not result in the upregulation of the transcription factor Tox. However, in line with the notion that T cell exhaustion is driven by TCR stimulation (Wherry et al., 2007; Mueller & Ahmed, 2009; Martinez et al., 2015), Tox was upregulated upon antigen-specific stimulation compared to cytokine stimulation in the PD-1 ⁺compartment (Fig. EV1F). Thus, while cytokines drive an improved effector response, they do not appear to exacerbate T cell exhaustion.

**Figure 1:**
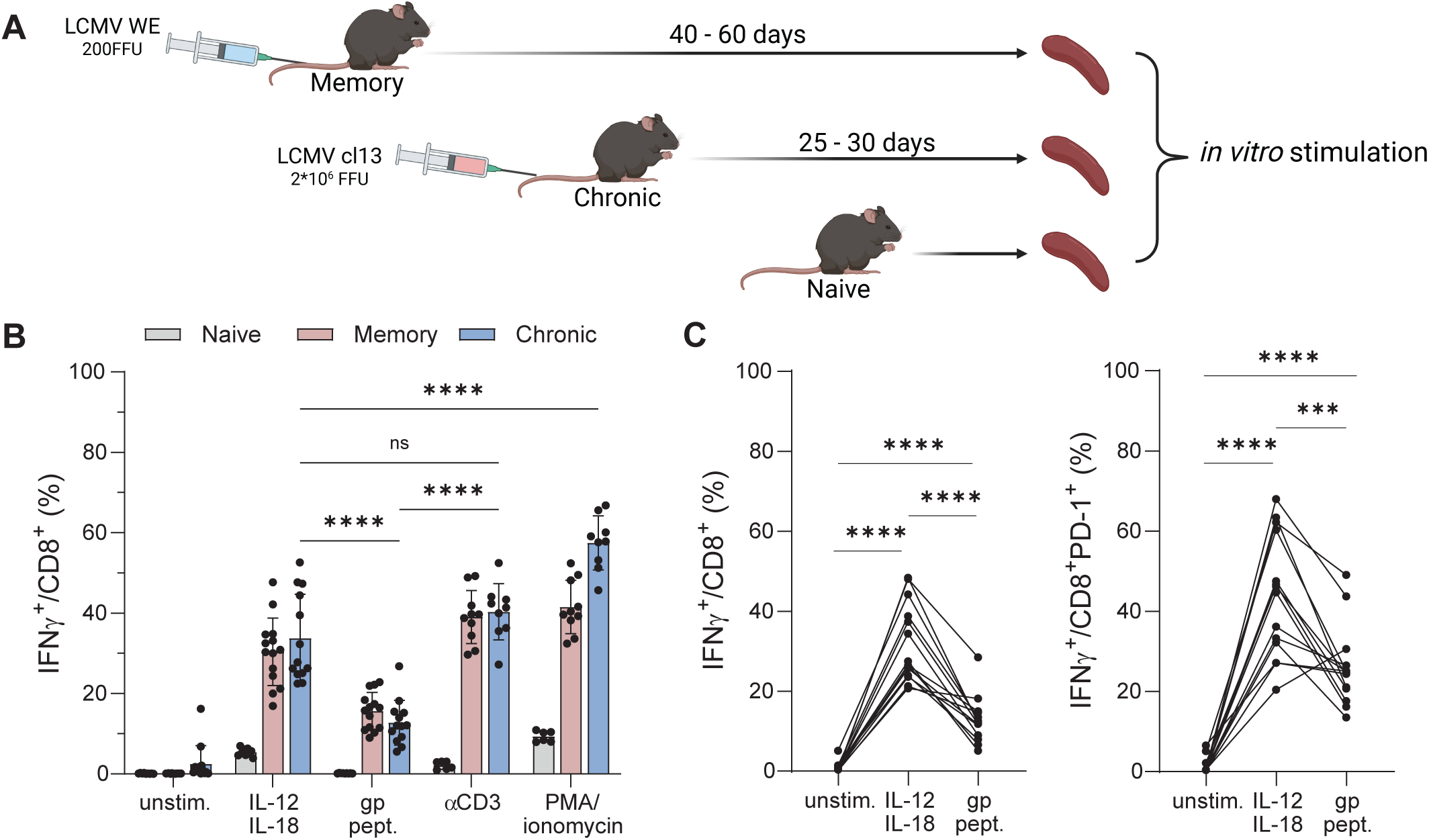
Exhausted T cells respond to cytokine stimulation. (**A**) Experimental setup of the in vitro assay using models of memory infection (LCMV WE d40-d60), chronic infection (LCMV Cl13 d25-d30) and naive mice. (**B**) Effector response (IFNg^+^) of CD8 T cells. (**C**) Paired analysis of the effector response between cytokine and gp peptide stimulated total CD8^+^ (left) or CD8^+^PD-1^+^ (right) cells from the spleen of mice with Cl13 infection. Results are pools of 2-3 experiments using 3-5 mice per group. Each dot represents one mouse; paired samples from the same mouse are connected by lines. Differences between groups were determined using two-way ANOVA (**B**), paired t test

Prior work concluded that exhausted CD8 ⁺ T cells in LCMV Cl13 infection are unresponsive to IL-12 and IL-18 (Ingram et al, 2011). To investigate this discrepancy, we examined IL - 18R expression. At baseline (unstimulated), IL - 18R levels in total CD8 ⁺ T cells from chronically infected mice were similar to those of memory CD8 ⁺ T cells; while the PD - 1 ⁺ CD8 ⁺ T cell subset showed reduced levels, IL-18R expression remained high (Fig. EV2A). Notably, stimulation with IL-12 and IL-18, but not with gp peptides, led to an upregulation of IL - 18R expression in both total as well as PD - 1 ⁺ CD8 ⁺ T cells (Fig. EV2B), suggesting that cytokine exposure itself enhances IL - 18R expression and potentially amplifies IL-18 responsiveness. Consistent with our previous studies on bystander T cells (Rakebrandt et al, 2024), we used an overnight stimulation protocol, while Ingram et al. applied only a short (5.5 h) cytokine stimulation. Given that we observe a continuous increase in IFNγ production in cytokine stimulated memory CD8^+^ T cells over time, with the peak in the range of 12-20h (Fig. EV2C), the shorter stimulation periods may not have captured the full potential of exhausted T cells to respond to cytokine stimulation.

### 3.2. Transcriptional signatures of innate-acting T cells can be found in exhausted cells

Innate-acting CD4^+^ T cell subsets (CD4^+^ T_IA_ cells) capable of responding to cytokine stimulation in a TCR-independent fashion show distinct transcriptional profiles (Rakebrandt et al, 2024). To define the transcriptional signature of bystander or innate-acting CD8^+^ T cells present in vivo, we generated a single-cell RNA-sequencing (scRNA-seq) dataset of CD8^+^ T cells isolated from a heterologous challenge model (Rakebrandt et al, 2024). In this model, LCMV-specific memory T cells are established through LCMV WE infection. After 40-60 days, when virus has been cleared and T cells memory has formed, mice are challenged with Legionella pneumophila (Lpn), which does not share antigens with LCMV and thus induces TCR-independent (bystander) activation and IFNγ production in a subset of innate-acting CD4^+^ and CD8^+^ T cells (T_IA_ cells) (Rakebrandt et al, 2024). CD8 ⁺ T cells from the lungs of LCMV WE memory and control mice (no previous infection) were sorted 2 days after Lpn challenge, enriched for the CD8^+^CD44^+^ compartment and subjected to scRNA-seq. We identified 12 CD8 ⁺ T cell clusters within this dataset (Fig. EV3A) that clearly segregated into naive and memory CD8 ⁺ T cells based on expression of Sell (CD62L) and Cd44 (CD44) (Fig. EV3B). Effector/memory cells (clusters 1, 2, 4, 5, 6, and 10, T_IA_ dataset) were re-clustered and categorized as Central Memory, Circulating/Proliferating Effector Memory (TEM), Effector-Memory, Terminal Effector, Interferon-stimulated (ISG), Trm-like and Innate-acting based on expression of canonical lineage, activation and functional markers (Fig. 2A and Fig. EV4A-C). We defined the Ifng expressing cluster (cluster 6) as Innate-acting T cells, as IFNγ production is their hallmark (Fig. 2C).

**Figure 2:**
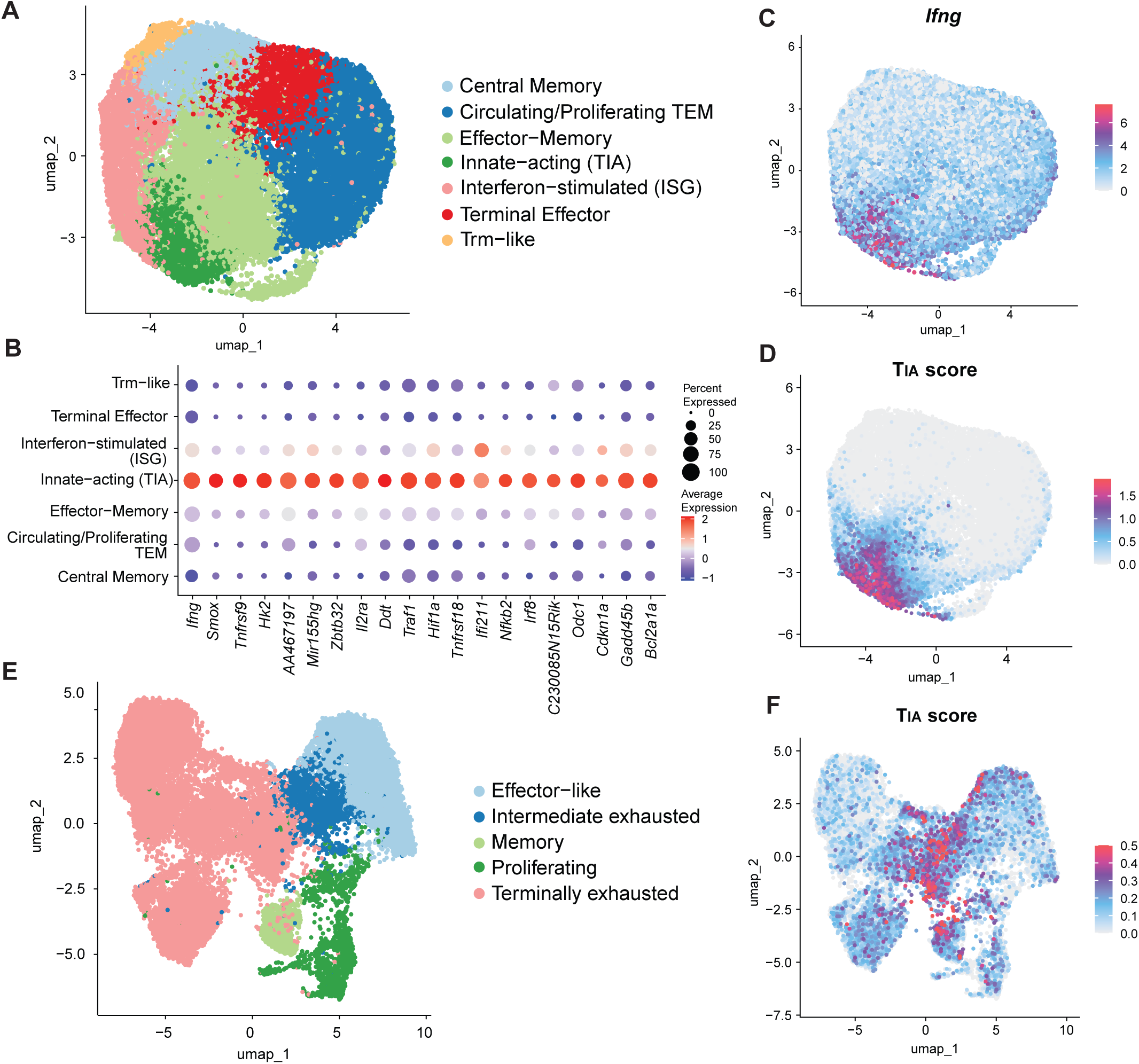
T_IA_ transcriptional signature is present in exhausted CD8^+^ T cells from Cl13 infection. (**A**) UMAP of scRNA-seq data of CD8^+^CD44^+^ T cells isolated from the lung of LCMV WE memory mice at d2 post Lpn infection. (**B**) Dot plot showing expression of 20 T_IA_ signature genes within this dataset. (**C** - **D**) Feature plot depicting *Ifng* expression (**C**) and T_IA_ score (**D**) within this dataset. (**E** - **F**) UMAP of scRNA-seq data of LCMV-specific P14 CD8^+^ T cells isolated on d21 of LCMV Cl13 infection (Sandu et al., 2020) depicting T cell subsets (**E**) and T_IA_ score (**F**) within this dataset.

To investigate the transcriptional similarities between T_IA_ cells and other T cell populations, we next defined a T_IA_ score based on the top 20 differentially expressed genes (DEGs) of the T_IA_ cluster compared to all other clusters in this dataset. To validate this signature, we calculated and projected the T_IA_ score onto our dataset, which showed strong enrichment specifically within the cluster identified as innate-acting CD8⁺ T cells (Fig. 2C, D). This score was then applied to public datasets of exhausted CD8^+^ T cells, to assess the presence of T_IA_-like transcriptional programs within exhausted T cell populations. We first examined a scRNA-seq dataset of LCMV-specific P14 CD8 ⁺ T cells from Cl13-infected mice (Sandu et al, 2020), collected at day 21 post infection (Fig. 2E). The T_IA_ score was applied to the dataset and mapped onto the UMAP (Fig. 2F). The T_IA_ signature was detected in discrete regions of the UMAP, primarily corresponding to terminally exhausted and intermediate exhausted cells, particularly at the interface of these two populations (Fig. 2E, F). Thus, although the T_IA_ signature does not correspond to a distinct exhausted CD8 ⁺ T cells subset, it marks a fraction of exhausted CD8 ⁺ T cells that emerge during chronic Cl13 infection.

### 3.3. Transcriptional and functional features of T_IA_ cells are found within tumor-infiltrating lymphocytes

Building on this finding, we next investigated whether the transcriptional and functional features of T_IA_ cells were also evident in other exhausted settings. To this end, we analyzed tumor-infiltrating lymphocytes (TILs), where T cell exhaustion is a dominant feature, to determine whether T_IA_-like properties represent a broader hallmark of exhausted CD8 ⁺ T cells across different contexts. Using the published scRNA-seq dataset from Mazet et. Al (2023), we restricted our analysis to TILs from WT mice while retaining the original cell-state annotations and UMAP embedding. We then applied the T_IA_ signature score to the WT subset and examined its distribution across stem-like, exhausted, and memory-bystander-like cells (Fig. 3A, B). The T_IA_ signature was predominantly enriched in the exhausted cells and, to a lesser extent, in the memory-bystander-like population, but was absent in the stem-like cells. These findings confirm that T_IA_ cells share transcriptional similarities with exhausted T cells present within tumors.

**Figure 3:**
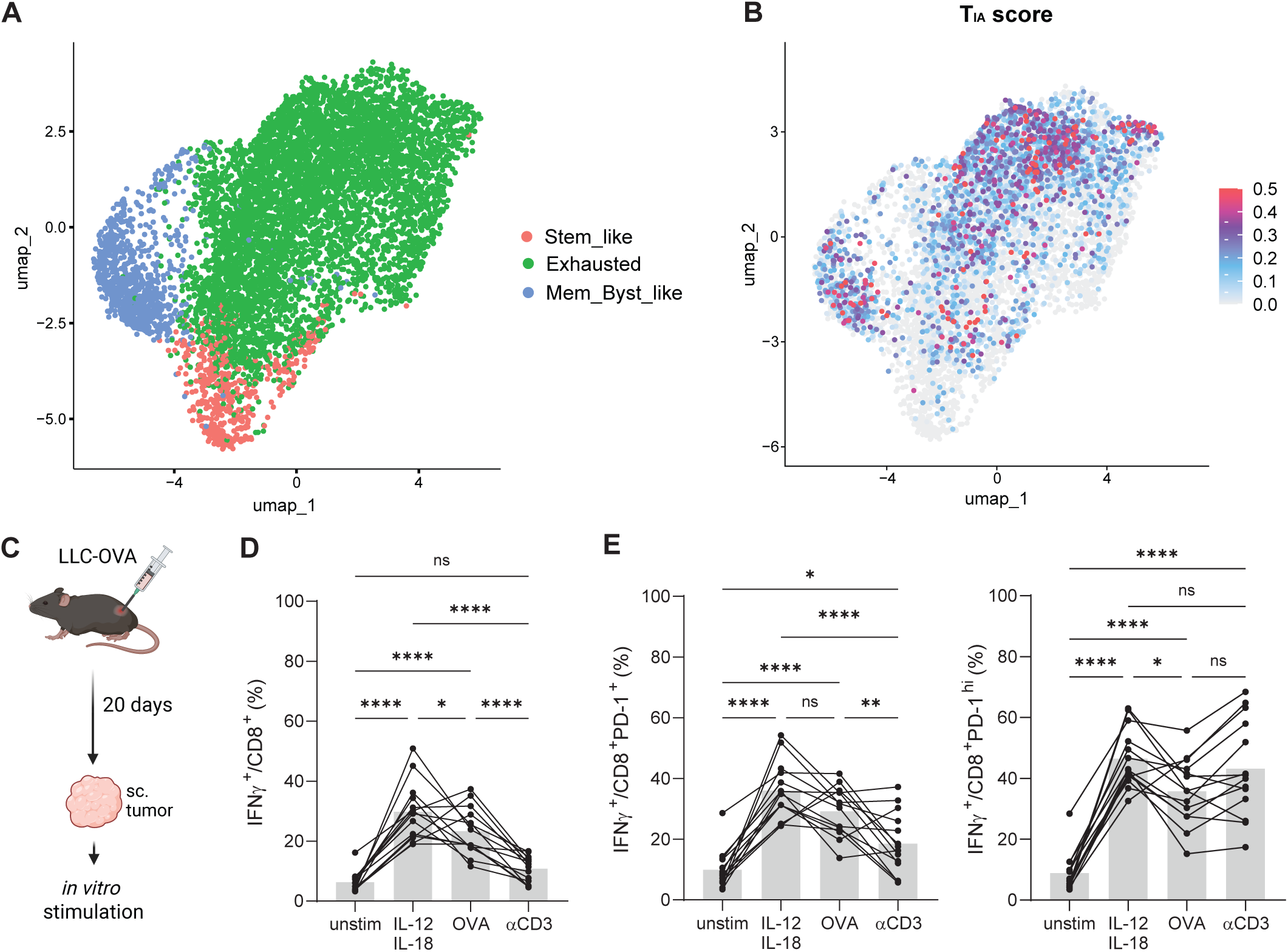
T_IA_ transcriptional signature and functional phenotype can be observed in cancer. (**A**) UMAP of WT tumor infiltrating CD8^+^ T cells (TILs) from the published scRNA-seq dataset of B16 tumors from Mazet et al., (2023) retaining original UMAP embedding and cell-state annotations. (**B**) Feature plot depicting TIA score within this dataset. (**C**) Experimental setup for results depicted in (**D**) and (**E**) TILs were isolated from LLC-OVA tumor bearing mice at d20 post cell transfer and stimulated in vitro. (**D** - **E**) TILs were stimulated with IL-12+IL-18, OVA peptide (OVA), anti-CD3 (αCD3), or left unstimulated (unstim) and IFNγ production within total CD8^+^ T cells (**D**) and PD-1^+^ (left) or PD-1^hi^ (right) CD8^+^ T cells (**E**) isolated from subcutaneous tumors was analyzed by flow cytometry. Results are pools of 2 experiments using 5-6 mice per group. Each dot represents one mouse; paired samples from the same mouse are connected by lines. Differences between groups were determined using two-way ANOVA (**D** and **E**). p < 0.05 (*), p < 0.01 (**), p < 0.001 (***).

Having observed these transcriptional similarities, we next assessed functional characteristics. To allow for assessment of antigen-specific responses, mice were injected with OVA-expressing Lewis Lung Carcinoma (LLC-OVA) cells and subcutaneous tumors were analyzed 20 days post induction (Fig. 3C). TILs were stimulated either in a TCR-dependent manner using αCD3 or OVA peptides or with IL-12 and IL-18 (Fig. 3D, E). In line with our observation in exhausted CD8^+^ T cells obtained from LCMV Cl13 infected mice, the IFNγ response following cytokine stimulation was higher than that observed following antigen-specific stimulation with OVA peptide for both total CD8^+^ T cells as well as for PD-1^hi^ exhausted CD8 ⁺ T cells (Fig. 3D, E). The enhanced overall response was primarily driven by PD-1^+^ cells (Fig. EV5), highlighting the ability of exhausted T cells to be activated in a bystander manner within the tumor microenvironment. Collectively, these data demonstrate that T_IA_-like transcriptional and functional features are shared by exhausted CD8+ T cells across diverse contexts, including cancer.

### 3.4. Bystander-activated exhausted T cells are protective in heterologous infectious challenge

We next explored whether this activation also occurs in vivo and influences subsequent immune responses. To this end, we infected mice with LCMV Cl13 to induce CD8^+^ T cell exhaustion and then challenged them during the chronic phase of the infection (day 25-30) with Legionella pneumophila (Lpn), a respiratory bacterial pathogen known to trigger the production of IL-12 and IL-18 (Spörri et al, 2006; Brieland et al, 2000). Importantly, IFNγ acts as the key protective cytokine in this infection model (Spörri et al, 2006). Mice were sacrificed during the innate phase of the Lpn infection (day 2), and lungs were harvested for analysis. We observed an early CD8 ⁺ T cells effector response marked by IFNγ production in the chronic group, but not in control mice (Fig. 4A). Interestingly, this response originated from both PD-1 ⁺ and PD-1 ⁻ cells (Fig. 4B), indicating that both exhausted and non-exhausted CD8^+^ T cells can respond in an innate-like manner during heterologous infection and are the dominant IFNγ producers at this stage of infection (Fig. 4C). We further examined the bacterial burden at this early time point and observed a significantly reduced bacterial titer in both the bronchoalveolar lavage (BAL) and lung tissue of LCMV Cl13 infected mice compared to Lpn challenges controls (Fig. 4D). Collectively these data suggest that bystander activation of exhausted T cells can also be observed in vivo and translates into enhanced pathogen control following heterologous bacterial challenge.

**Figure 4:**
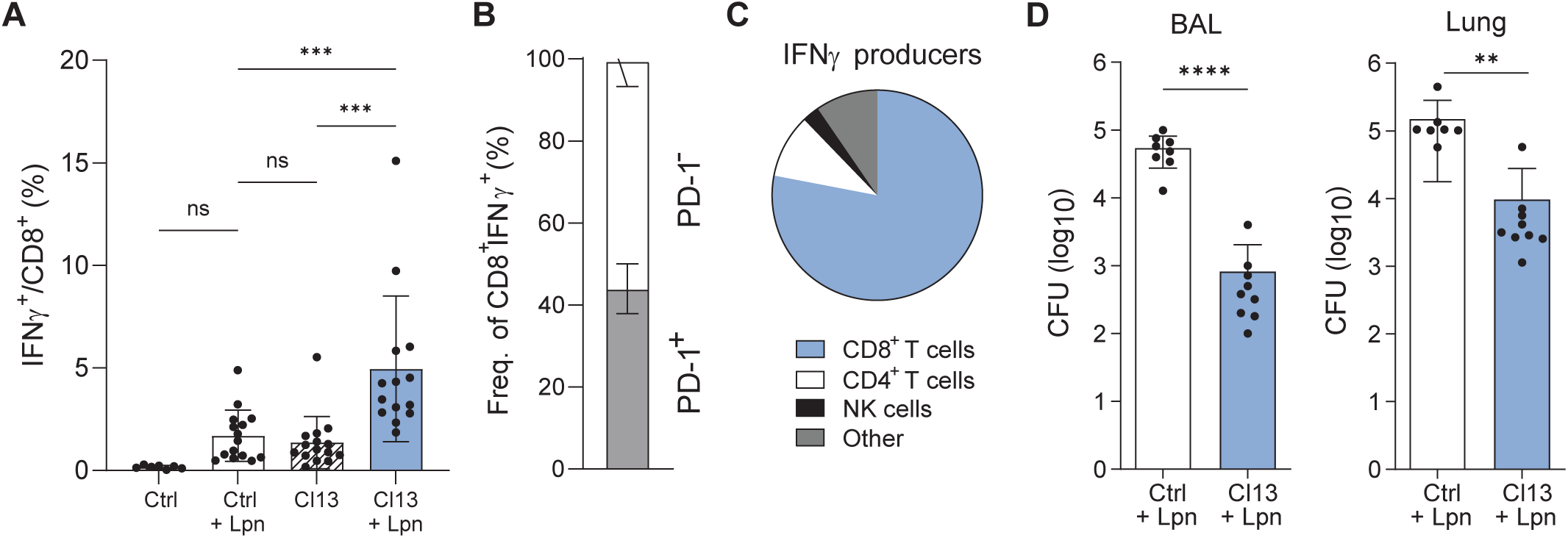
Cytokine activated exhausted cells contribute to protection from heterologous challenge. Mice were chronically infected with LCMV Cl13 (Cl13) or left naive as controls (Ctrl) and challenged with Lpn i.n. on day 25-30. (**A**) IFNγ response in CD8^+^ T cells in the lung on d2 post Lpn infection. (**B**) Distribution of PD-1^+^ vs PD-1^-^ CD8^+^ T cells within the IFNγ+ CD8^+^ T cell population. (**C**) Proportion of CD8^+^ T cells, CD4^+^ T cells, NK cells, and other populations among total live IFNγ^+^ cells. (**D**) Lpn CFU 60h p.i. in BAL (left) and lung (right). Pooled data of 2-3 experiments using 4-5 mice per group. Differences between groups were determined using one-way ANOVA (A), multiple unpaired t test (C) and unpaired t test (D). p < 0.05 (*), p < 0.01 (**), p < 0.001 (***) and p < 0.0001 (****).

## 4. DISCUSSION

CD8^+^ T cell exhaustion is a hallmark of chronic infection and cancer and defined by sustained antigen exposure, high expression of inhibitory receptors, and impaired effector functions (McLane et al, 2019; Wherry et al, 2007). In both chronic viral infections and tumors, exhaustion is viewed as a largely irreversible state, with terminally differentiated cells exhibiting poor proliferative potential and limited effector function (Miller et al, 2019; Siddiqui et al, 2019; Utzschneider et al, 2016). Memory CD8 ⁺ T cells are known to be able to undergo antigen-independent bystander activation in response to innate cytokines such as IL-12 and IL-18 (Freeman et al, 2012; Raué et al, 2012; Soudja et al, 2012). Here, we show that exhausted CD8 ⁺ T cells also possess the capacity for bystander activation via IL-12 and IL-18, enabling effector responses comparable to or exceeding antigen-specific stimulation, both in vitro and in vivo.

Our findings reveal that cytokine-induced activation of PD-1 ⁺CD8^+^ cells is not accompanied by upregulation of Tox, a key driver of the exhaustion program (Khan et al, 2019; Alfei et al, 2019). Furthermore, we found that IL-12 and IL-18 stimulation itself enhanced IL-18R expression, and that extended cytokine stimulation provided sufficient time for receptor upregulation and subsequent effector responses. This provides a potential explanation for the discrepancy of our results with prior studies concluding unresponsiveness of exhausted T cells to cytokine stimulation, which tested much shorter stimulation conditions (Ingram et al, 2011). Taken together our results suggest that IL-12 and IL-18 stimulation engages activation pathways distinct from those triggered by antigen exposure, allowing exhausted CD8^+^ T cells to mount functional responses. Rather than being universally unresponsive, exhausted CD8 ⁺ T cells retain inducible effector potential, revealing an unexpected flexibility in their functional capacity.

We also identified transcriptional similarities between these bystander-responsive exhausted CD8 ⁺ T cells and a distinct population of innate-acting CD8 ⁺ T_IA_ cells. Applying the newly generated transcriptional T_IA_ signature to public datasets from chronic LCMV infection (Sandu et al, 2020) and murine tumors (Mazet et al, 2023), we detected T_IA_-like profiles within exhausted CD8 ⁺ T cell populations, particularly at the interface between intermediate and terminal exhaustion. This suggests that exhausted cells can reside in a repressed state regarding antigen-dependent activation but remain responsive to cytokine stimulation, allowing for rapid mobilization of effector potential under the right inflammatory signals. This is particularly relevant in the tumor setting as terminally exhausted TILs are poorly responsive to antigen and checkpoint blockade (Siddiqui et al, 2019; Miller et al, 2019). We found that terminally exhausted PD-1^hi^ CD8^+^ T cells still responded to cytokine stimulation, suggesting that while these cells may be less responsive to TCR-dependent signals, they can still be mobilized through cytokine-mediated activation. Importantly, we show that cytokine responsiveness of exhausted CD8^+^ T cells is not limited to in vitro culture conditions but also takes place in vivo, highlighting the physiological relevance of our findings. Moreover, we have previously demonstrated in CD4^+^ T cells that cytokine responsiveness may be more ubiquitous than previously appreciated, extending beyond conventional memory subsets (Yassini et al, 2025). This raises the possibility of therapeutic strategies combining cytokines like IL-12 and IL-18 with checkpoint blockade to enhance antitumor immunity, consistent with prior reports of IL-12 based interventions in cancer models (Tugues et al, 2014).

In conclusion, our results demonstrate that exhausted CD8 ⁺ T cells can be reactivated through IL-12 and IL-18 stimulation, exhibiting both functional and transcriptional features shared with innate-acting T cells. These findings challenge the classical view of exhaustion as a fixed, irreversible state and suggest that innate cytokine pathways represent an alternative route to reinvigorate exhausted T cell responses in chronic infection and cancer.

## 5. MATERIALS AND METHODS

### 5.1. Mice

C57BL/6 (B6) mice were purchased from Janvier Labs. housed in individually ventilated cages containing autoclaved bedding and nesting material, with standard diet and water ad libitum on a 12-h light/12-h dark cycle (18–23 °C, 40–60% humidity) at LASC Zürich, Switzerland.

### 5.2. Ethical statement

All experiments were performed in accordance with institutional policies and regulations of the relevant animal welfare acts and have been reviewed and approved by the Cantonal veterinary office in Zurich, Switzerland.

### 5.3. Viral and Bacterial Infections

For the acute LCMV WE and the chronic LCMV Cl13 infections, 200 FFU and 2×10^6^ FFU, of the respective strains, were injected i.v.. For L. pneumophila (Lpn) infections, 3×10^6^ CFU of Lpn JR32 FlaA^−^ (Weber et al, 2012) was applied intranasally. 40 to 60 day after the primary acute LCMV WE infection or 25 to 30 days after chronic LCMV Cl13 infection, mice were infected with 3 × 10^6^ CFU Lpn JR32 FlaA^−^ (Weber et al., 2012), grown on charcoal yeast agar plates. At sacrifice, animals were perfused with PBS and bronchoalveolar lavage (BAL), lungs, and/or spleen were collected. For determination of bacterial titers, BAL and lungs from infected mice were collected, lungs were lysed using a Qiagen Tissue Lyser II, and samples were plated on charcoal yeast agar plates and grown for 3 d at 37 °C.

### 5.4. In vitro Assays

Spleens were processed by mechanical disruption in RPMI 1640 medium supplemented with 10% FCS, penicillin (100 IU/mL), and 1% L-glutamine. Red blood cells were lysed with ACK buffer (155 mM NH4Cl, 10 mM KHCO3, and 0.1 mM Na2EDTA, pH: 7.4) for 3 min. For stimulations, single cell suspensions were incubated in U-bottom plates with the indicated cytokines (10 ng/mL, BioLegend) 16-20h overnight with 5 μg/mL Brefeldin A (BioLegend) for the last 4h; re-stimulated with gp_33-41_ (gp33, 1ug/ml, KAVYNFATC, EMC microcolletions GmbH) and gp_61-80_ (GLKGPDIYKGVYQFKSVEFD, 1ug/ml, EMC microcolletions GmbH) for 4 hours, or with anti-CD3 (2 µg/mL, clone 145-2C11; BioXCell) for 4 hours, or left unstimulated. All incubations were performed at 37°C and 10% CO_2_.

### 5.5. Tumor models

For tumor cell inoculation, the murine lung cancer cell line Lewis lung carcinoma (LLC) expressing ovalbumin (LLC-OVA) (Herbst et al, 2025), was cultured in DMEM (Gibco) containing 10% FCS (Capricorn), 100 U/mL penicillin, 100 μg/mL streptomycin (Gibco) and 2 mM L-glutamine (Gibco) and 5 x 10^5^ LLC-OVA cells were injected subcutaneously into the right flank. After 20 days, tumors were harvested in RPMI 1640 containing above mentioned supplements, cut into small pieces using scissors and digested using 1 mg/mL Collagenase Type IV (Gibco) and 0.2 mg/mL DNase I (Roche). After 45 min incubation at 37°C under continuous rotation, digestion was stopped using ice cold supplemented RPMI and cell suspensions were filtered through 70 μm cell strainers followed by an ACK lysis (2 mL, 1 min). Isolated immune cells were restimulated with 2 μg/mL OVA peptides (OVA_257-264_ and OVA_323-339_, EMC) or 1 ug/mL anti-CD3 (clone 145-2C11, BioXCell) in the presence of 5 μg/mL Brefeldin A (BioLegend) for 4h at 37°C.

### 5.6. Flow Cytometry

FACS stainings were performed on single-cell suspensions from spleen and lung. Spleen was processed as described above. Lungs were enzymatically digested with collagenase D (Gibco) and DNase I (VWR) for 30 min, and immune cells were isolated using a 30% Percoll (GE Healthcare) gradient as described previously (Rakebrandt et al, 2024). When staining for intracellular cytokines, cells were incubated with Brefeldin A (BioLegend) for 4 h at 37 °C prior to staining. For surface staining, antibodies were incubated for 20 to 30 min at RT in PBS. The Zombie NIR fixable dye (BioLegend) or LIVE/DEAD fixable Blue dye (Invitrogen) was used to exclude dead cells and debris. For intracellular cytokine staining, cells were permeabilized using the Cytofix/Cytoperm kit (BD Biosciences) for 5 to 8 min at RT, followed by antibody incubation for 20 to 30 min at RT.

The following antibodies were used for flow cytometry: αCD4-BUV496 (RM4-5), αCD4-FITC (RM4-5), αCD8a-BUV395 (53-6.7), αCD44-AlexaFluor700 (IM7), αPD-1-BUV737 (RMP1-30) and αNK1.1-BUV615 (PK136) were purchased from BD., αCD8a-SuperBright600 (53-6.7), αIL-18Ra-PerCP/eFluor710 (P3TUNYA), Rat IgG2a-PE (eBR2a) purchased from eBioscience. αTox-PE (TXRX10) was purchased from Invitrogen. αCD44-PerCP (IM7), αIFNγ-PE/Dazzle594 (XMG1.2) and high concentration streptavidin-APC were all purchased from Biolegend. Tetramer was prepared in-house using gp33-biotin and concentrated streptavidin-APC.

Data were acquired on a Cytek Aurora 5L and analyzed using SpectroFlo (Cytek) and FlowJo software (TreeStar). The gating strategies used to identify the respective T-cell populations are shown in Fig. EV6.

### 5.7. Single-cell RNAseq

#### 5.7.1. Sample preparation and sequencing

Lung single-cell suspensions were obtained from 6 LCMV-WE memory mice (d40-60 p.i.) challenged with Lpn (d2 p.i.). Both infections and lung processing were done as described above. At single cell suspension stage, cells from two mice were pooled to obtain sufficient cell numbers. Cells were stained and sorted for CD8^+^CD4^-^CD44^hi^ cells, resuspended and incubated in 50ul FACS buffer with 0.5ul of TrueStrain FcX^TM^ PLUS (anti-mouse TruStain FcX™ PLUS (anti-mouse CD16/32) CD16/32) blocking agent (Biolegend) for 10 minutes at 4°C. Each sorted sample was then labeled using anti-mouse Hashtag reagents (TotalSeq B0301, B0302 and B0303, BioLegend) before pooling. Library preparation was performed by the Functional Genomic Center Zurich using 10x Chromium Next GEM Single Cell 5’ Reagent kits, according to manufacturer’s instructions (10x Genomics). scRNA-seq/TCR-seq libraries were sequenced on an Illumina Novaseq 6000 (Illumina).

#### 5.7.2. Analysis and data visualization

The raw sequences in the fastq files were aligned to the mouse mm10 reference transcriptome using CellRanger (4.0.0). The empty droplets from each sample were removed using emptyDrops (Lun et al, 2019) from the DropletUtils package (Heiser et al, 2021), and doublets were removed using Scrublet (Wolock et al, 2019).

The gene expression data was then normalized using the SCTransform (Hafemeister & Satija) function from Seurat (Satija et al, 2015). Louvain community clustering (Satija et al, 2015) was used to find cell clusters. Upon cluster definition their identity was confirmed by analyzing expression of Cd3e and CD8a, and two small, separated clusters, which did not express these two markers were removed. Data was then re-normalized, re-clustered and further analyzed in R 4.4.1 (R Core Team, 2025) using the Seurat package (Hao et al, 2021). The packages tidyverse (Wickham et al, 2019), ggplot2 (Wickham, 2016), and dplyr (Wickham et al, 2023) were used for visualizations.

#### 5.7.3. Annotation of the T_IA_ dataset

To assign phenotypic identities to transcriptionally defined cell clusters in the TI_A_ dataset, we performed manual annotation guided by the expression of established lineage, activation and effector-associated markers. Marker genes were selected based on well-described roles in CD8^+^ T cell differentiation, migration and activation (Chen et al, 2021; Sandu et al, 2020; Kuhn et al, 2022; van der Gracht et al, 2021; Kaech & Wherry, 2007; Milner et al, 2020; Kaech & Cui, 2012; Jia et al, 2023).

Memory-associated and recirculating populations were characterized by expressions of Klf2, S1pr1, Tcf7, Sell, Ccr7, Cxcr3 and Il7r. Effector and cytotoxic phenotypes were defined by upregulation of Tbx21, Runx3, Cx3cr1, Gzmb, Prf1, Klrk1 and Ccr5, reflecting terminal effector differentiation and cytolytic function. Mk67 expression was used to determine potential proliferative populations. Activation signatures were captured by Cd69, Il2ra (CD25), Ifng, and Il18r1 were used as markers for responsiveness to inflammatory cues, while a distinct interferon responsive population was defined by enrichment of interferon-stimulated genes (ISG). Several clusters exhibited overlapping effector, homing and persistence markers and occupied intermediate positions between memory and effector states in the UMAP space.

Given their shared transcriptional features and lack of clear separation, these clusters were consolidated and annotated as ‘intermediate’. Based on the gene expression and cluster similarity analysis, clusters were annotated as Central Memory, Circulating/Proliferating Effector Memory (TEM), Effector-Memory, Terminal Effector, Interferon-stimulated (ISG), Trm-like and Innate-acting or TIA (Fig. 2A and Fig. EV4).

#### 5.7.4. T_IA_ transcriptional signature

The cluster containing T_IA_ cells was identified based on expression of Ifng, which represents the primary functional marker of the cell population of interest. Following cluster identification, differential gene expression analysis was performed between the TIA cluster and all remaining cells in our dataset using the FindMarkers function in Seurat V5.4.0 (Satija et al, 2015). Resulting genes were filtered for statistical significance (adjusted p value < 0.05) and biological relevance (−1< average log2 fold change < 1). Genes were subsequently ranked by average log2 fold change in descending order and further filtered based on expression prevalence, requiring expression in at least 25% of cells (pct > 0.25). The top 20 genes meeting these criteria were defined as the transcriptional signature of T_IA_ cells.

#### 5.7.5. Publicly available scRNAseq data

The publicly available single-cell RNA-seq datasets were kindly provided by the authors as processed Seurat objects of publicly available single-cell RNA-seq datasets (Mazet et al., 2023; Sandu et al., 2020). Data of P14 cells (Sandu et al, 2020) were reprojected in UMAP, but the original subset annotations were retained (Fig. 2E). Data was further analyzed in R v4.5.2 (R Core Team, 2025) using the Seurat package v5.4.0 (Hao et al, 2021).

### 5.8. Statistics

All statistical analyses were performed using GraphPad Prism and were two-sided. Data was tested for normality with Shapiro-Wilk test and QQ-plot analysis. All results are shown as mean ± SD. Statistical significance is defined as p<0.05 and shown as *, p<0.01 as **, p<0.001 as ***, and p<0.0001 as ****.

## 6. DATA AVAILABILITY

The scRNA-seq data from CD8^+^ T cells has been deposited on the ArrayExpress database at EMBL-EBI (www.ebi.ac.uk/arrayexpress) and is available via accession number E-MTAB-16869. Publicly available datasets used in this study include the scRNA-seq dataset from GEO (GSE221118) and raw sequencing data from the European Nucleotide Archive (ENA; PRJEB36998). Processed data (Seurat object) provided by the original authors were used where available.

## 7. AUTHOR CONTRIBUTIONS

Conceptualization: E.G. and N.J. Experimentation: E.G., R.D., N.Y., N.R., and Analysis: E.G., R.D. Formal Analysis: E.G. and Z.L. Writing – Original Draft: E.G. and N.J. Review and Editing: all authors. Funding Acquisition: N.J. Supervision: F.J.Q. and N.J.

## 9. ACKNOWLEDGEMENTS

We would like to thank the Joller and Oxenius group members for helpful discussions. This work was supported by the Swiss National Science Foundation (PP00P3_181037, 310030_197590 to N.J.) and the European Research Council (677200 Immune Regulation to N.J.).

## EXPANDED VIEW FIGURE LEGENDS

**Fig. EV1: Exhaustion state is not altered by cytokine stimulation**

CD8^+^ T cells from Cl13 infected mice (d25-30) were analyzed under different in vitro conditions. Unstimulated - unst., gp pepstides - gp pept. (**A**) Representative FACS plots of PD-1 and Tox expression in unstimulated CD8^+^ T cells isolated from naive mice (left) or mice with chronic (middle) vs. acute (right) LCMV infection. (**B**) IFNγ production and expression of IL18Rα in tetramer^+^ PD-1^hi^ cells upon cytokine stimulation. (**C**) Expression of Tox in tetramer^+^ PD-1^hi^ cells under unstimulated conditions. (**D**) Representative FACS plots of IFNγ response of CD8^+^ T cells from an acute LCMV (left) and chronic LCMV infection (right) upon gp restimulation. (**E**) Frequency of PD-1^hi^ and PD-1^int^ cells within total CD8^+^ T cells following different in vitro stimulations. (**F**) Tox expression in CD8^+^PD-1^+^ (left) or total CD8^+^ (right) T cells was assessed post in vitro assay by flow cytometry. Results are pools of 1 (**C, F**). 2 (**B**) or 3 (**E**) experiments using 4-5 mice per group. Each dot represents one mouse; paired samples from the same mouse are connected by lines. Differences between groups were determined using two-way ANOVA (**E**) and paired t test (**F**). p < 0.05 (*).

**Fig. EV2: CD8^+^ T cells from a chronic infection upregulate IL18Rα upon prolonged cytokine exposure.**

Splenic CD8^+^ T cells from LCMV memory mice (LCMV WE d40-d60), mice chronically infected with (LCMV Cl13 d25-d30) on naïve controls were analyzed by flow cytometry. (**A**) IL18Rα expression in total (left) or PD-1^+^ (right) CD8^+^ T cells. (**B**) IL18Rα expression in total (top) or PD-1^+^ (bottom) CD8^+^ T cells from a chronic infection after in vitro stimulation with IL-12 and IL-18, LCMV gp peptides or in unstimulated controls. (**C**) IFNγ production by purified splenic CD8^+^ T cells from naïve and LCMV memory mice following in vitro IL-12 and IL-18 stimulation over time. Pooled data of 3 (**A, B**) or 2 (**C**) experiments using 2-5 mice per group. Each dot represents one mouse; paired samples from the same mouse are connected by lines. Differences between groups were determined using two-way ANOVA (**A, C**) and paired t test (**B**). p < 0.05 (*), p < 0.01 (**), p < 0.001 (***) and p < 0.0001 (****).

**Fig. EV3: CD8 T cells from murine lungs.**

(A) UMAP oaf Seurat clusters of scRNA-Seq data from total CD8+ T cells from lungs of control (top) and LCMV memory (bottom) mice at d2 of Lpn infection. (**B**) Violin plot indicating expression of Sell (top) and Cd44 (bottom) in the Seurat clusters.

**Fig. EV4: Characterization of T_IA_ scRNA-Seq dataset.**

(**A**) UMAP of T_IA_ dataset. (**B**) Dot plot of marker genes used for cluster annotation. (**C**) Heatmap of average gene expression per cluster with genes used for annotation of T_IA_ dataset and the cluster similarity based on the canonical markers.

**Fig. EV5: PD-1 expression in CD8^+^ T cells responding to cytokine stimulation.**

TILs were isolated from LLC-OVA tumor bearing mice at d20 post cell transfer and stimulated in vitro with IL-12 and IL-18 or OVA peptide (OVA) and (**A**) IFNγ production within PD-1^+^ and PD-1^-^ CD8^+^ T cells or (**B**) PD-1 expression within IFNγ^+^ CD8^+^ T cells was analyzed by flow cytometry. Pooled data of 2 experiments using 5-6 mice per group.

**Fig. EV6: Gating strategy and representative flow cytometry plots.**

(**A**.) Representative gating strategy for CD8^+^ T cells and assessment of PD-1, tetramer (tet), and IL-18R expression. (**B**.) Representative plot of Tox gating within PD-1^+^ CD8^+^ T cells from Cl13.

