## Supplementary Figures for "Cytokine-mediated activation of exhausted T cells"

### *Expanded View Figures*

### Figure EV1

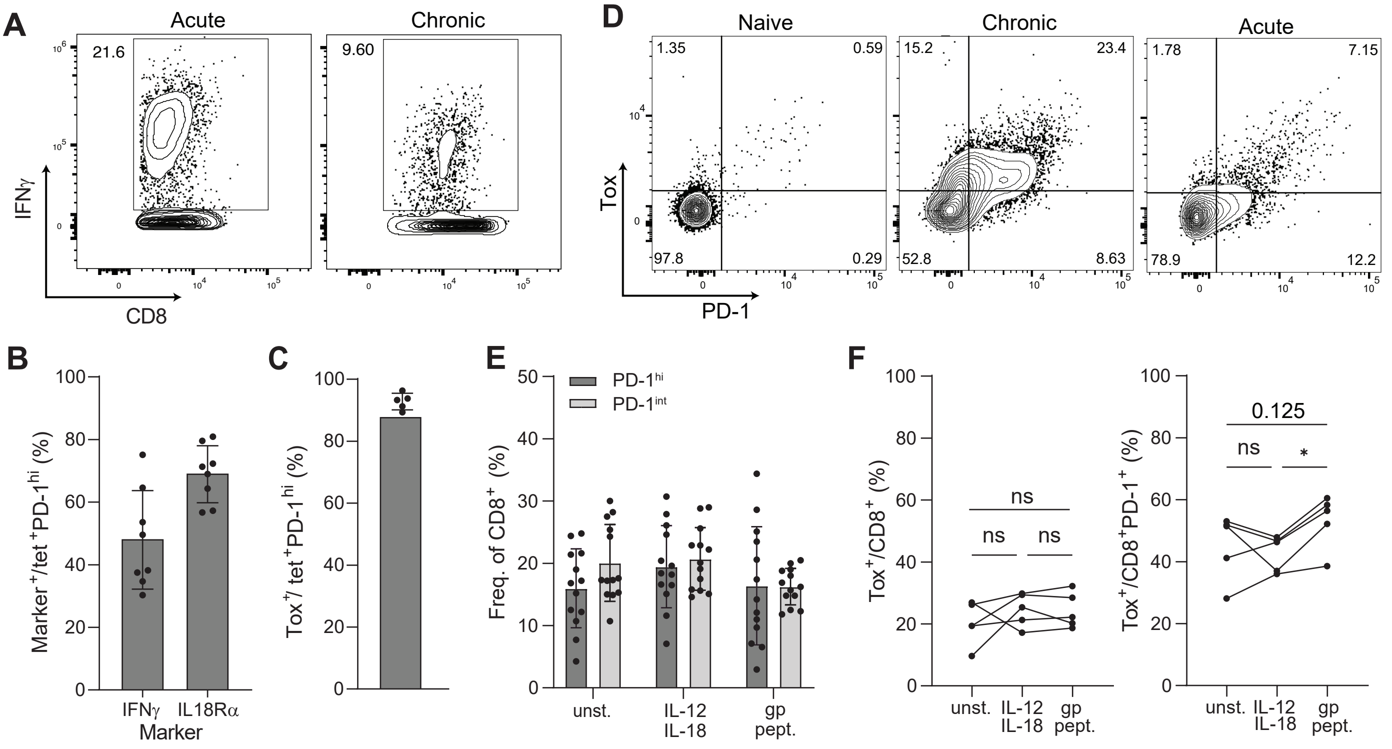


**Figure EV1: Exhaustion state is not altered by cytokine stimulation**

CD8⁺ T cells from Cl13 infected mice (d25-30) were analyzed under different in vitro conditions. Unstimulated - unst., gp pepstides - gp pept. (**A**) Representative FACS plots of PD-1 and Tox expression in unstimulated CD8⁺ T cells isolated from naive mice (left) or mice with chronic (middle) vs. acute (right) LCMV infection. (**B**) IFNγ production and expression of IL18Rα in tetramer⁺ PD-1$^{hi}$ cells upon cytokine stimulation. (**C**) Expression of Tox in tetramer⁺ PD-1$^{hi}$ cells under unstimulated conditions. (**D**) Representative FACS plots of IFNγ response of CD8⁺ T cells from an acute LCMV (left) and chronic LCMV infection (right) upon gp restimulation. (**E**) Frequency of PD-1$^{hi}$ and PD-1$^{int}$ cells within total CD8⁺ T cells following different in vitro stimulations. (**F**) Tox expression in CD8⁺PD-1⁺ (left) or total CD8⁺ (right) T cells was assessed post in vitro assay by flow cytometry. Results are pools of 1 (**C, F**). 2 (**B**) or 3 (**E**) experiments using 4-5 mice per group. Each dot represents one mouse; paired samples from the same mouse are connected by lines. Differences between groups were determined using two-way ANOVA (**E**) and paired t test (**F**). $p < 0.05$ (*).

#### Figure EV2

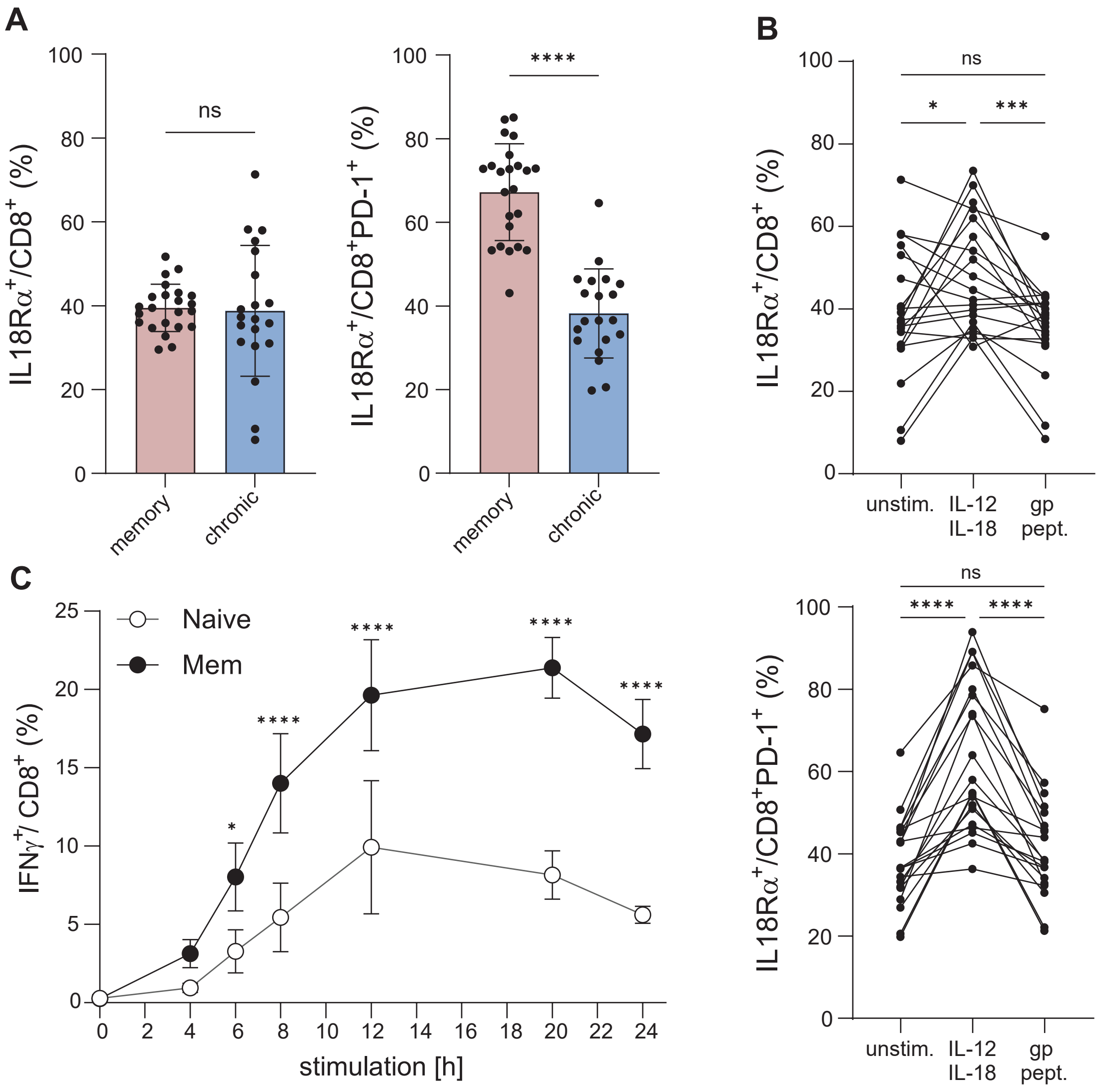


**Figure EV2: CD8⁺ T cells from a chronic infection upregulate IL18Rα upon prolonged cytokine exposure.**

Splenic CD8⁺ T cells from LCMV memory mice (LCMV WE d40-d60), mice chronically infected with (LCMV Cl13 d25-d30) on naïve mice (Control) were analyzed by flow cytometry. (**A**) IL18Rα expression in total (left) or PD-1⁺ (right) CD8⁺ T cells. (**B**) IL18Rα expression in total (top) or PD-1⁺ (bottom) CD8⁺ T cells from a chronic infection after in vitro stimulation with IL-12 and IL-18, LCMV gp peptides or in unstimulated controls. (**C**) IFNγ production by purified splenic CD8⁺ T cells from naive and LCMV memory mice following in vitro IL-12 and IL-18 stimulation over time. Pooled data of 3 (**A, B**) or 2 (C) experiments using 2-5 mice per group. Each dot represents one mouse; paired samples from the same mouse are connected by lines.Differences between groups were determined using two-way ANOVA (**A, C**) and paired t test (**B**). $p < 0.05$ (*), $p < 0.01$ (**), $p < 0.001$ (***) and $p < 0.0001$ (****).

#### Figure EV3

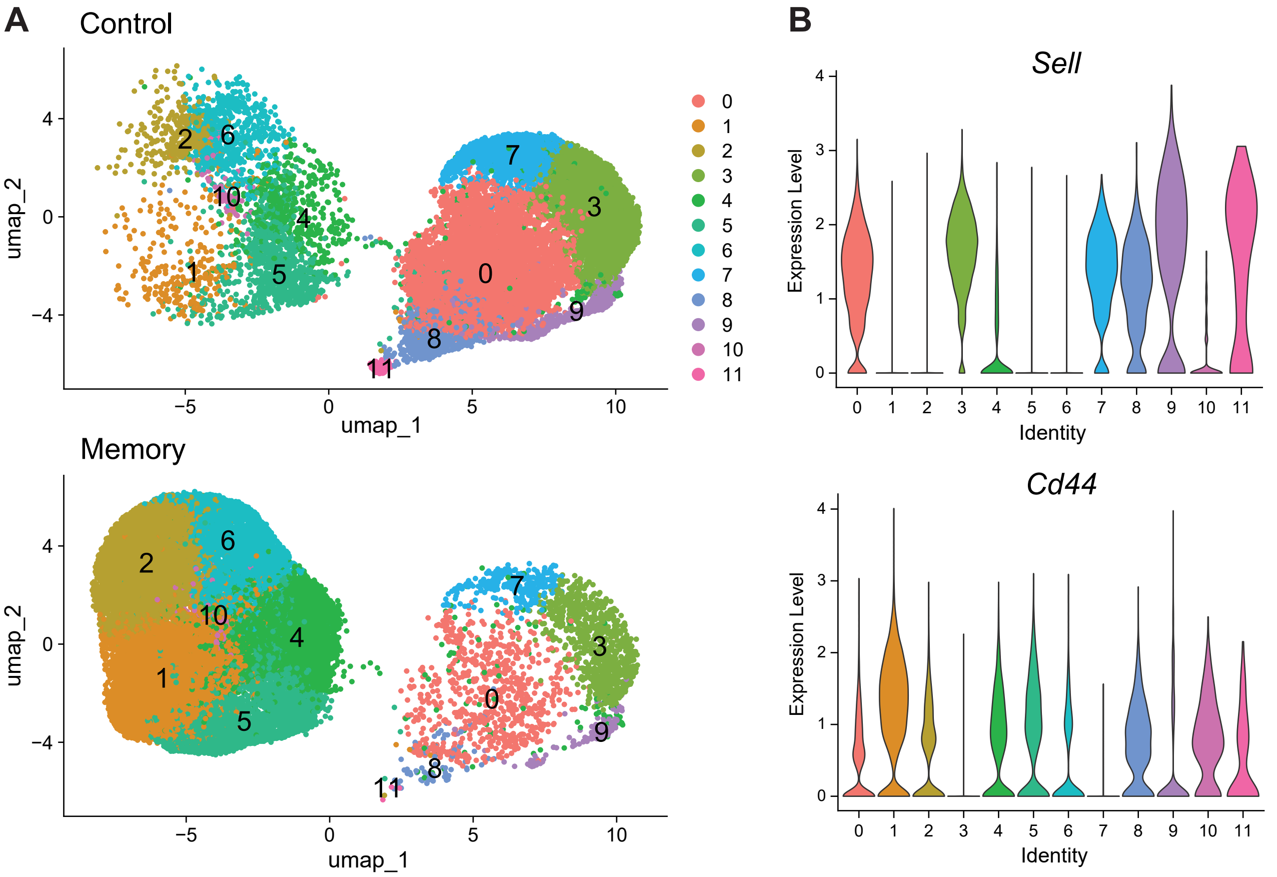


**Figure EV3: CD8 T cells from murine lungs.**
(**A**) UMAP of Seurat clusters of scRNA-Seq data from total CD8⁺ T cells from lungs of control (top) and LCMV memory (bottom) mice at d2 of Lpn infection. (**B**) Violin plot indicating expression of *Sell* (top) and *Cd44* (bottom) in the Seurat clusters.

#### Figure EV4

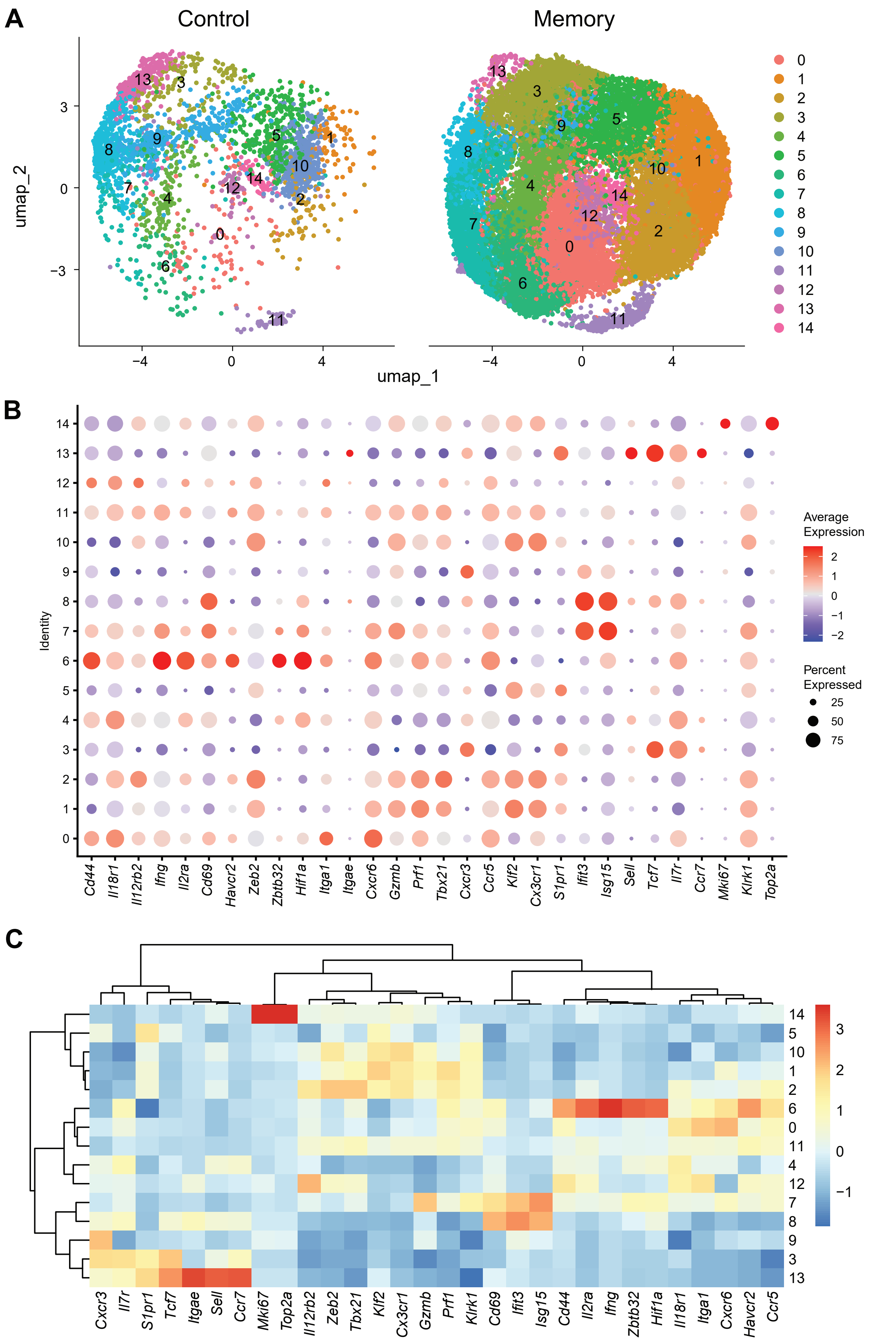


**Figure EV4: Characterization of $T_{IA}$ scRNA-Seq dataset.**
(**A**) UMAP of $T_{IA}$ dataset. (**B**) Dot plot of marker genes used for cluster annotation. (**C**) Heatmap of average gene expression per cluster with genes used for annotation of $T_{IA}$ dataset and the cluster similarity based on the canonical markers.

### Figure EV5

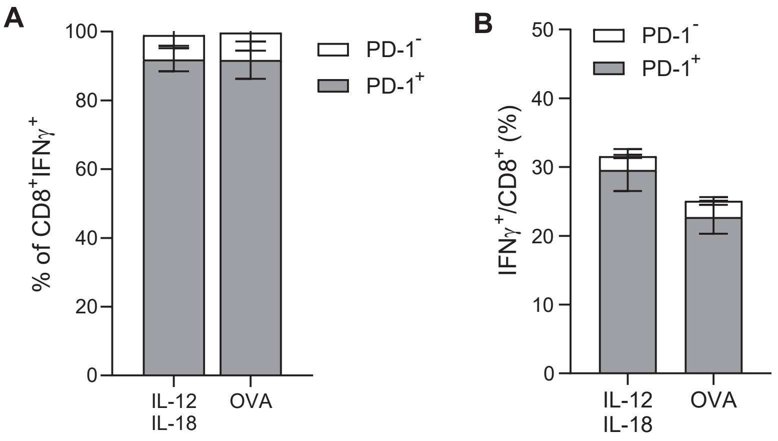


**Figure EV5: PD-1 expression in $CD8^+$ T cells responding to cytokine stimulation.**
TILs were isolated from LLC-OVA tumor bearing mice at d20 post cell transfer and stimulated *in vitro* with IL-12 and IL-18 or OVA peptide (OVA) and (**A**.) IFNγ production within $PD\text{-}1^+$ and $PD\text{-}1^-$ $CD8^+$ T cells or (**B**.) PD-1 expression within $IFN\gamma^+$ $CD8^+$ T cells was analyzed by flow cytometry. Pooled data of 2 experiments using 5-6 mice per group.

### Figure EV6

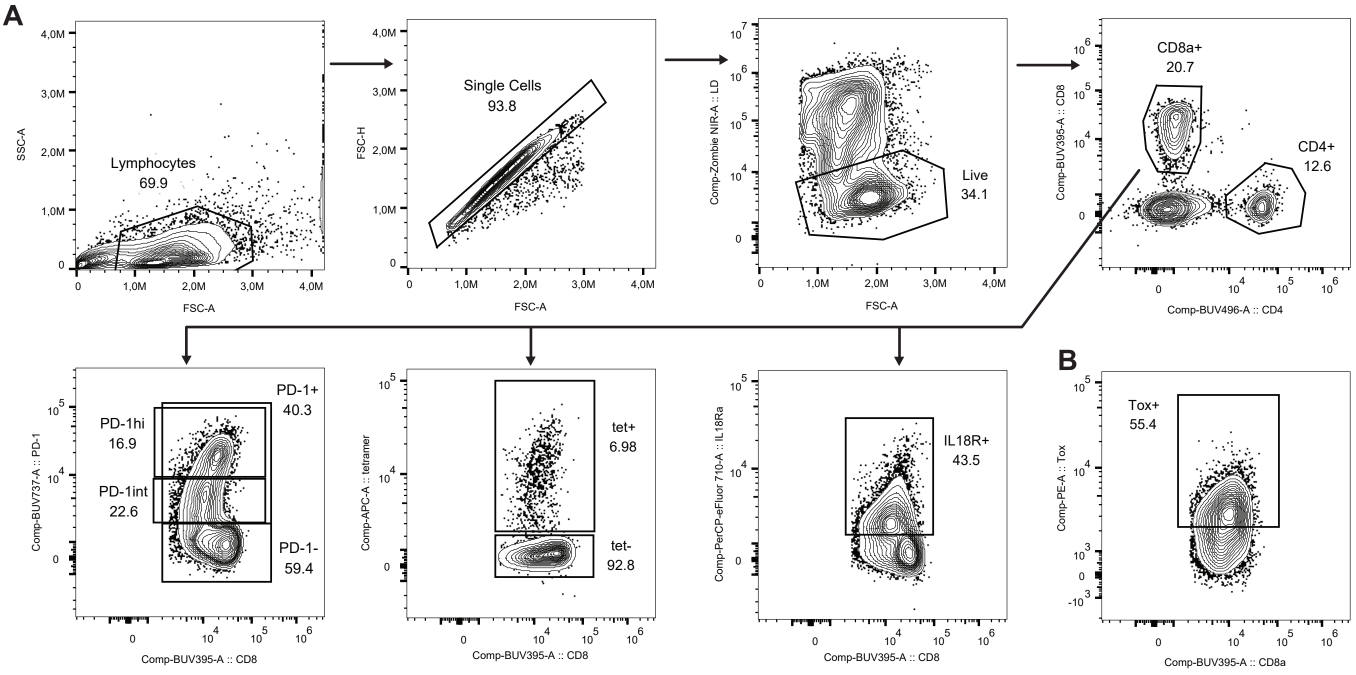


**Figure EV6: Gating strategy and representative flow cytometry plots.**
(**A**.) Representative gating strategy for CD8⁺ T cells and assessment of PD-1, tetramer (tet), and IL-18R expression. (**B**.) Representative plot of Tox gating within PD-1⁺ CD8⁺ T cells.
